# Threshold inclusion size triggers conversion of huntingtin to prion-like state that is reversible in newly born cells

**DOI:** 10.1101/2023.02.13.528394

**Authors:** Leila Asgarkhani, Ibrahim Khandakar, Rachel Pakan, Theresa C. Swayne, Lesley Emtage

## Abstract

Aggregation of mutant Huntingtin protein (mHtt) leads to neuronal cell death and human disease. We investigated the effect of inclusion formation on yeast cells. Previous work indicates that mHtt protein moves both in and out of inclusions, potentially undergoing refolding in the inclusion. However, the sustained influx of unfolded protein into an inclusion leads to a dramatic change from a phase-separated body to an irregular, less soluble form at a threshold inclusion size. Altered morphology was associated with a prion-like seeding that accelerated inclusion growth despite loss of soluble cytoplasmic protein. The structural change abolished exchange of material between the inclusion and the cytosol and resulted in early cell death. Affected cells continued to divide occasionally, giving rise to daughters with a similar phenotype. Most newly born cells were able to reverse the prion-like aggregation, restoring both soluble cytoplasmic protein and a normal inclusion structure.

## INTRODUCTION

Destabilizing mutations in an abundant protein can result in an abnormally high and continuous burden of unfolded protein leading to aggregation, as is the case with familial neurodegenerative diseases. In Huntington’s disease, the expansion of a polyglutamine (polyQ)-encoding tract in HTT exon 1 results in the release and aggregation of an N-terminal fragment of the Htt protein.^1,2^ Using *S. cerevisiae*, we have expressed mutant Huntingtin exon 1 fused to GFP (mHtt^ex1^-GFP) from a strong constitutive promoter. As in mammalian cells, inclusions of mutant Htt^ex1^-GFP form in wild-type *S. cerevisiae* with a frequency proportional to the length of the polyQ tract.^3,4^

While micron-sized aggregates of mHtt^ex1^ will form spontaneously in solution^5–8^, living cells control the aggregation process. Yeast cells lacking either the chaperone Hsp104 or the low complexity protein Rnq1 form neither inclusions nor visible small particles of mutant mHtt^ex1^- GFP.^3,9^ That is, mHtt^ex1^-GFP aggregates large enough to be visible by high-resolution light microscopy do not form spontaneously in yeast cells. Instead, the cellular machinery promotes inclusion formation, and cellular factors are found in the inclusion. Most mHtt^ex1^-GFP inclusions in yeast are ovoid condensates that concentrate Hsp104 together with unfolded mHtt^ex1^-GFP.^10^

Although ovoid inclusion bodies have gel-like physical properties and are overwhelmingly singular, their functional qualities are likely similar to those of stress-induced condensates which also contain Hsp104 and concentrated unfolded proteins. Hsp104 is known to act as a disaggregase in a concentration-dependent manner,^11^ and Q-bodies and stationary phase granules have both been shown to process and release processed protein in an Hsp104-dependent manner and will typically diminish and disappear within a few hours of stress cessation.^12–14^

However, unlike transient stresses that induce Q-bodies and stationary-phase granules, the high expression of unstable proteins results in a continuous flow of unfolded protein into an inclusion that may exceed the capacity of the inclusion to process it.

Since intracellular inclusions were first identified in brain autopsies over 100 years ago, and as ever more intracellular phase-separated structures have been discovered, there has been uncertainty about their roles and consequences. Some inclusions, like Q-bodies, are clearly adaptive; others, such as those in the substantia nigra of Huntington’s patients, are strongly associated with cell death. Here, we studied the fate of mHtt^ex1^-GFP inclusions and of the cells containing them, using time-lapse imaging to track cells for several hours and across several divisions. We identified a series of structural changes concomitant with a change in the balance between diffuse and inclusion-bound protein, with the late stages in this process associated with prion-like seeding and cell death. Strikingly, daughter cells born at this stage were typically able to recover a normal phenotype, demonstrating a physiological mechanism for reversing the prion-like state.

## RESULTS

While most mHtt^ex1^-GFP inclusions in actively growing cells are mobile and ovoid, a minority of cells contain large, unmoving asymmetrical inclusions, often in combination with small inclusions dispersed throughout the cell. Even more rarely, cells contain small, dispersed inclusions but no large inclusion. In mammalian cells, Ramdzan and colleagues^15^ have shown that the contents of large Htt-GFP inclusions undergo structural shifts from a more to a less accessible form. To better understand relationships between the variety of inclusions, we followed the growth and development of inclusions in individual cells using time-lapse imaging.

### Inclusion structure changes predictably over time

Cells were transformed with either mHtt^ex1^(72Q)-GFP or mHtt^ex1^(103Q)-GFP. Log-phase cells were imaged every hour for 6-9 hours; imaging frequency was limited to reduce photobleaching and photodamage. We followed the transformation and growth of large inclusions in 87 cells (52 cells expressing 103Q, 35 expressing 72Q).

Inclusions underwent predictable, stereotyped changes over time: the distribution of material within the inclusion became increasingly non-uniform as the inclusion shape became more irregular (Fig. 1). We divided inclusion development into three stages: the initial ovoid stage, an intermediate stage, and a late stage marked by internal heterogeneity, which we have named clastic, because it is reminiscent of clastic sedimentary rocks formed from accretion of heterogeneous rock fragments.

**Figure 1.**
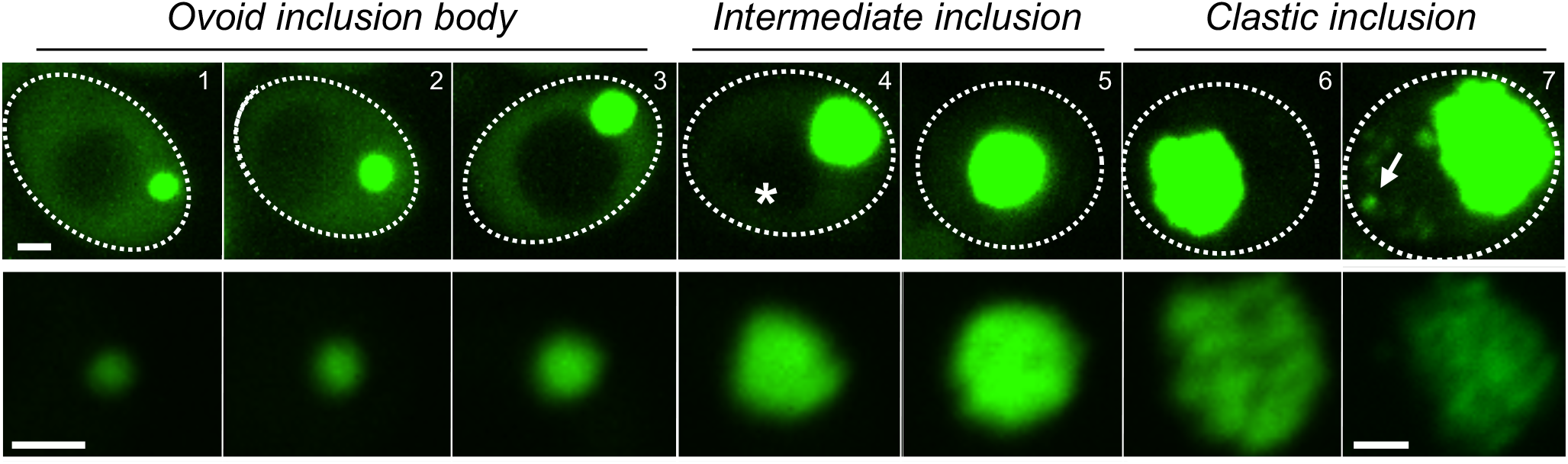
As ovoid inclusion bodies grow, they undergo a predictable set of visible structural changes. Upper row: representative images of cells showing characteristic changes in inclusion appearance over time. Lower row: magnified view of each inclusion shown in the upper image. Images have been contrast-adjusted to see the cytoplasm where possible (upper row), or the distribution of the inclusion contents (lower row). To encompass the full set of changes over time, three individual cells are shown: images 1-4 and 5-6 are of two different cells, images are 2-3 hours apart. Changes in inclusion structure are accompanied first by a marked reduction in the soluble cytoplasmic mHtt-GFP (*), then by the appearance of small intense inclusions dispersed throughout the cell (arrow). Images are of cells expressing mHtt^ex1^(72Q)-GFP but are equally characteristic of mHtt^ex1^(103Q)-GFP. Cell outlines indicated with dotted line; scale bars, 1 µm.

Expression of mHtt^ex1^-GFP in this system is constitutive, and therefore the cytoplasmic pool is being constantly replenished through protein synthesis, as well as by release from the ovoid inclusion.^10^ Transformation of inclusion structure coincided with a sharp drop in diffuse, soluble cytoplasmic mHtt^ex1^-GFP protein. Surprisingly, the near disappearance of cytoplasmic protein did not stop growth of inclusions. Eventually, small but intense inclusions appeared, dispersed throughout the cell (Fig. 1). Together, the abrupt loss of soluble protein and accumulation of aggregates are consistent with a prion-like seeding event.^16–18^ The complete structural transformation of the inclusion took place over a variable length of time, typically multiple cell cycles.

Our first goal was to develop an objective analytical method to classify inclusion stages and thereby identify the early stages of structural transition in both time-lapse and single images.

We found that visible heterogeneity of the contents of the inclusions was the earliest change that could consistently be recognized. Heterogeneities in the inclusion were identified by segmentation using a local thresholding algorithm (Supplementary Fig. 1a) on an optical section through the middle of the inclusion.^19–21^ Local thresholding detects variation in intensity within objects. Using this method, higher intensity regions are detected as objects, and internal lower intensity regions are detected as void space.

Ovoid inclusions show relatively homogenous intensity, and therefore have little or no void space, typically 0-2% (Supplementary Fig. 1b). Void area in intermediate inclusions rose abruptly over 2% and typically remained at a higher level (Supplementary Fig 1c). For this reason, void area was used as the criterion for determining the initial point of structural transition, while an increased number of locally thresholded objects was used to distinguish between intermediate and clastic inclusions. Inclusions were therefore categorized as ovoid (single, ≤ 2% void space), intermediate (single, >2% void space), or clastic (multiple objects).

### Structural changes and protein seeding are triggered by inclusion size

We identified 52 cells containing an inclusion that underwent a transition(s) between structural stages over the imaging timecourse. The most common transition was that of an ovoid inclusion body to an intermediate inclusion (n=35) or an intermediate to a clastic inclusion (n=24). Less often, ovoid inclusions appeared to transition directly to clastic inclusions (n=10). Some inclusions were observed in all 3 stages (n=17). An example of an inclusion that underwent both transitions is shown in Fig. 2a.

**Figure 2.**
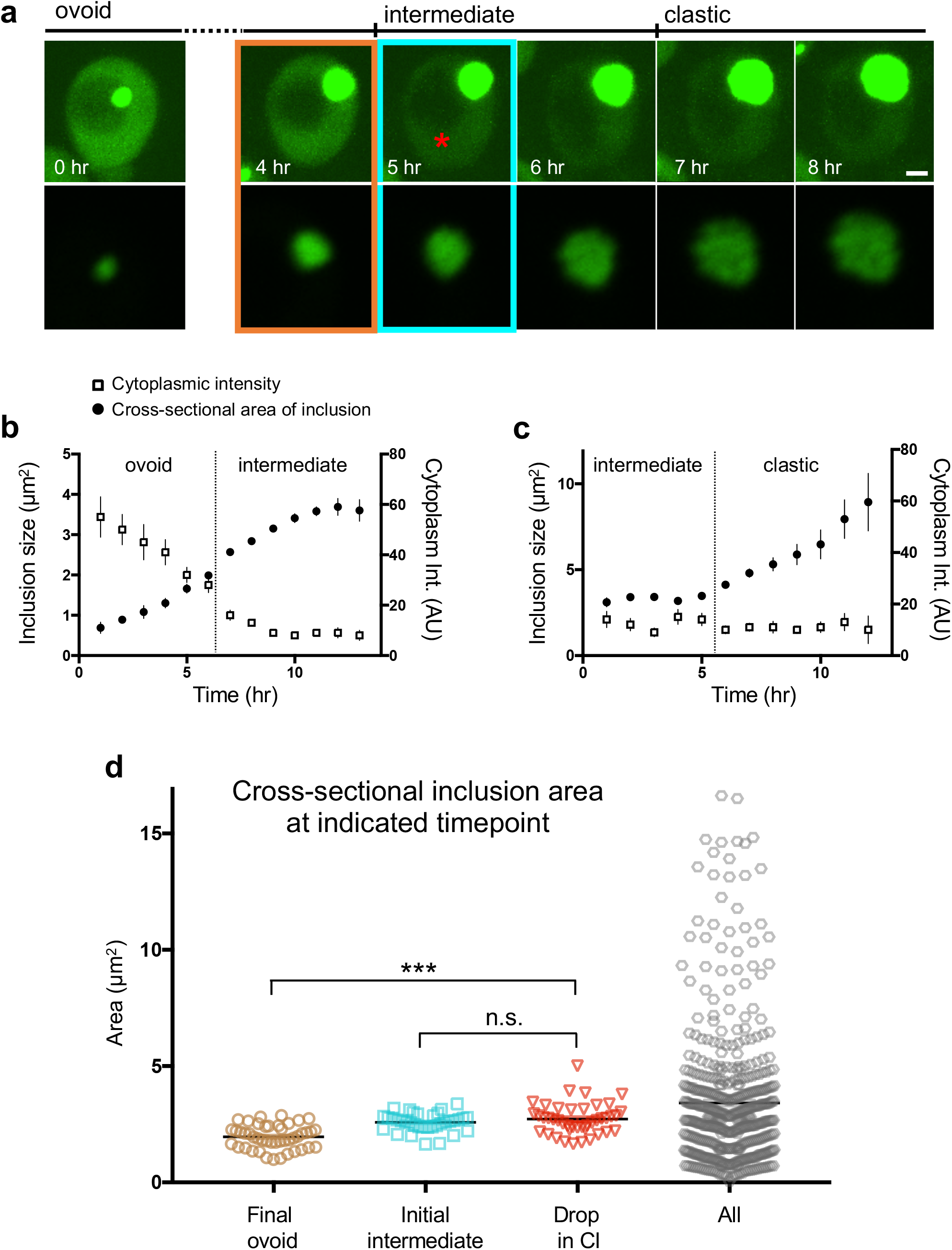
Structural transformation and loss of cytoplasmic mHtt are initiated coordinately. **a**, Images of an inclusion over 8 hours. Upper row: maximum intensity projection at indicated timepoints, contrast-adjusted to view the cytoplasm; lower row: magnified optical section of inclusion, contrast-adjusted to reveal the internal distribution of mHtt(72Q)-GFP. Orange and blue boxes highlight the transition from ovoid to intermediate, and the red asterisk indicates the initial timepoint with a substantial drop in cytoplasmic intensity. Scale bar, 1 μm. **b-c**, The cross-sectional area of mHtt-GFP inclusions undergoing the transition from ovoid to intermediate (**b**) or intermediate to clastic stage (**c**) is plotted with the cytoplasmic intensity of mHtt-GFP in the same cells over the same period (error bars indicate SEM, n=7-35 or 6-23 inclusions at each timepoint, respectively, combined 72Q and 103Q data sets). Vertical line indicates the point of transition. **d**, Cross-sectional inclusion area at critical transitions is shown: the final ovoid inclusion body timepoint (Final ovoid), the initial timepoint after the transition to the intermediate state (Initial intermediate), and the timepoint at which the first sharp drop in cytoplasmic intensity was observed (Drop in CI). Cross-sectional areas of inclusions from all timepoints are included for reference (All, n=495). The difference between the cross-sectional inclusion areas at the Final ovoid and the Drop in CI timepoints was highly significant, whereas areas at the Initial intermediate and Drop in CI points were not significantly different (p<0.0001 for Final ovoid to Drop in CI; p=0.77 for Initial intermediate to Drop in CI; ANOVA, Tukey’s multiple comparison test, n=45, 35, and 42. Line indicates mean, error bar indicates SEM).

To determine the relationship between inclusion size, cytoplasmic mHtt, and inclusion morphology, we measured the cytoplasmic intensity of mHtt-GFP and the cross-sectional area in the central focal plane of the inclusion for each cell at each timepoint, categorizing inclusion stage according to the criteria outlined above. To assess inclusion area and cytoplasmic mHtt-GFP levels at structural transition points, we aligned measurements at the first timepoint where the inclusion was identified as either intermediate or clastic. Fig. 2b shows the mean cytoplasmic mHtt-GFP intensity and cross-sectional area of inclusions that were observed to transition from ovoid to intermediate, aligned at the first intermediate timepoint. Fig. 2c shows the corresponding data for inclusions that transitioned from intermediate to clastic. The largest change in cytoplasmic intensity occurred at the ovoid-to-intermediate transition: an 85% decline.

Next, we asked whether the drop in cytoplasmic mHtt levels preceded the structural transformation of the inclusion, or whether the change in cytoplasmic levels and structural transition occurred concomitantly. Because inclusions grow continuously during this process, we used inclusion size as a “clock” to look for temporal correlations between structural transformation and cytoplasmic intensity. Fig. 2d shows the cross-sectional area of inclusions at the last timepoint in which they were categorized as ovoid, the first timepoint of the intermediate stage, and the first timepoint at which cytoplasmic mHtt-GFP levels dropped sharply. Inclusion size when the drop in cytoplasmic levels was first observed was significantly different from inclusion size immediately before the structural transition, but not from its size at the transition point (Fig. 2d). We conclude that there is an underlying process linking both the ovoid-to-intermediate transition and the loss of diffuse cytoplasmic protein.

We note that the areas of the 35 inclusions at the transition to the intermediate stage fell into a relatively narrow range (Fig. 2d). This suggests that the common process leading to the inclusion’s structural transformation and the depletion of the soluble cytoplasmic pool of mHtt-GFP may be initiated at a threshold inclusion size.

### Inclusion growth rate slows at the initiation of structural changes, then accelerates despite loss of soluble cytoplasmic protein

Measurements of cross-sectional area suggested that inclusion growth slowed temporarily as structural changes began, and then accelerated (Fig. 2b, c). We began by establishing the length of time that it took for inclusions to pass through the intermediate stage. To calculate an average value for the duration of the intermediate stage, we considered inclusions that were observed in all three stages, those that did not pass through an intermediate stage, and those that were observed for at least 3 hours in the intermediate stage, even if they did not transition during that time. The average length of time in the intermediate phase for these cells was 3.0 ± 0.4 hr (mean ± SEM; Supplementary Fig. 2). This value is likely to be an underestimate, because the full length of the intermediate stage was not observed for most inclusions in this data set.

To facilitate comparison of the kinetics of inclusion growth, inclusion volume was normalized to the size at the point of transition. Ovoid to intermediate transitions and intermediate to clastic transitions were analyzed separately (Fig. 3a–b). A typical timecourse for all stages was reconstructed by combining the datasets from both transitions assuming an intermediate stage duration of 4 hours (Fig. 3c) and normalizing to inclusion size at the ovoid-intermediate transition. Data from individual inclusions are shown in Supplementary Fig 3a, b. Our analysis shows that the inclusion growth rate slowed at the start of the intermediate stage, and the inclusion resumed exponential growth as it entered the clastic stage. The changes in growth rate coincide with the changes in structure, and therefore support the 3-stage classification we defined based on morphology.

**Figure 3.**
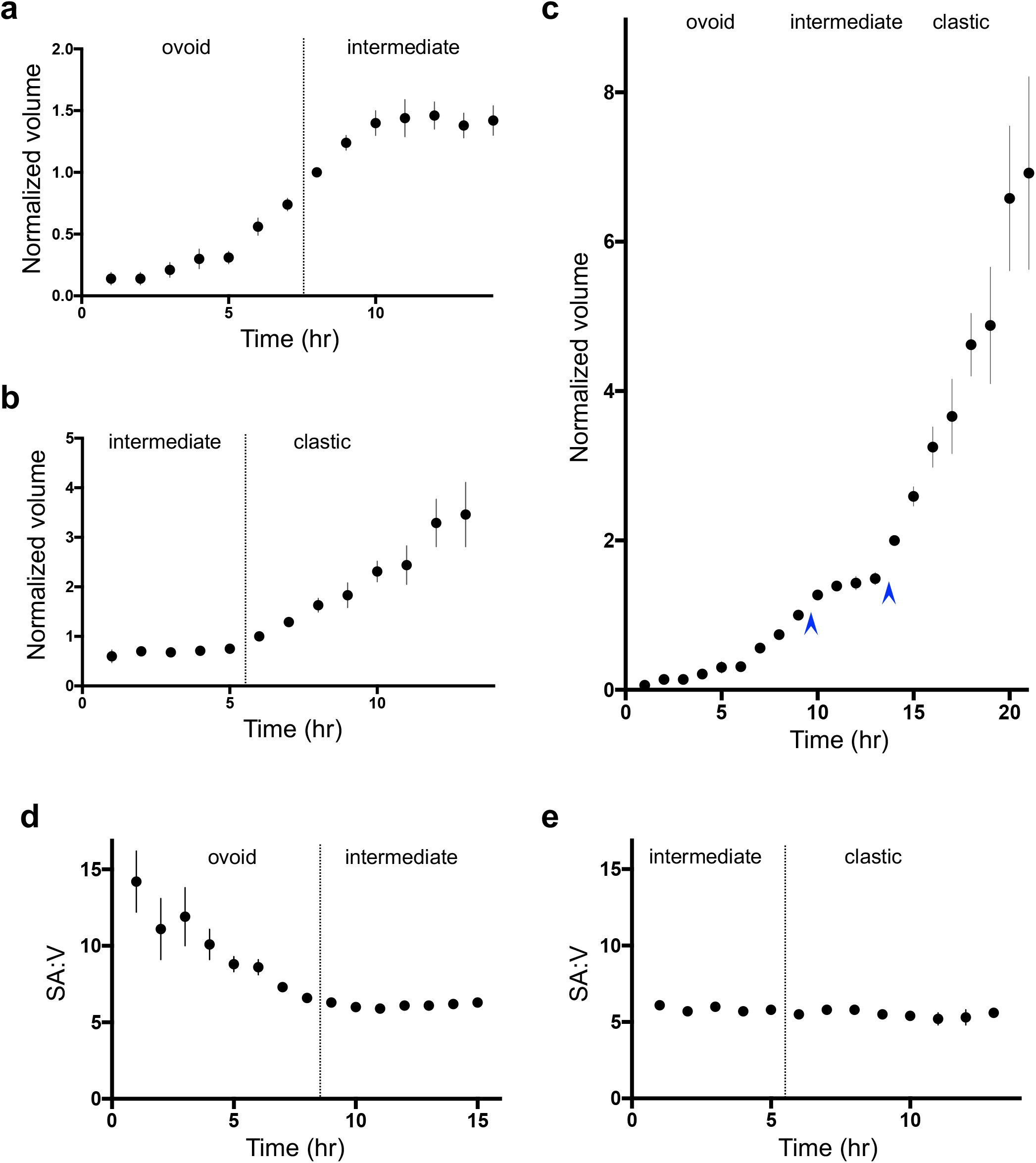
Inclusions undergo changes in growth rate associated with structural changes. **a**, Normalized volume of inclusions in cells expressing mHtt(72Q) and mHtt(103Q)-GFP as they enter the intermediate stage. Inclusion volume was normalized and aligned to that at the first intermediate stage (dashed line; error bars represent SEM, n=4-22 inclusions). **b**, Normalized volume of inclusions transitioning from intermediate to clastic stage. Inclusion volume was normalized and aligned to the first clastic-stage volume (dashed line); error bars represent SEM, n=2-20 inclusions per timepoint. **C**, Data from all timecourses combined, assuming intermediate stage duration of 4 hours and normalized to volume at the ovoid-intermediate transition (arrows indicate transitions; error bars represent SEM, n=2-22). **d, e**, Surface area-to-volume ratio of mHtt(72Q) and mHtt(103Q)-GFP inclusions. Mean values are shown for ovoid to intermediate and intermediate to clastic-stage inclusions (error bars represent SEM, n=3-19 and 2-18 inclusions, respectively).

During structural transitions, inclusions became increasingly irregular in shape. We used our measurements to calculate the surface area to volume ratio at the transition from ovoid to intermediate and intermediate to clastic inclusions (Fig 3d, e). The ratio of surface area to volume of a growing spherical inclusion is expected to decrease proportional to the reciprocal of the radius. For growing ovoid inclusions, surface area to volume decreased, consistent with a spherical or ellipsoid shape, but the ratio levelled out at the point of transition to the intermediate stage and did not change significantly again. These results demonstrate that the inclusion surface became increasingly convoluted as inclusion structure changed.

Remarkably, clastic stage inclusions continued to grow exponentially even after the cytoplasmic mHtt^ex1^-GFP fell to near background levels, demonstrating that new mHtt-GFP is being synthesized though not accumulating as soluble cytoplasmic protein. However, clastic inclusion volume did show a sigmoidal trend (Fig. 3c) and eventually reached a plateau (Supplementary Fig. 4a, b).

Cytoplasmic mHtt^ex1^-GFP levels did not significantly decline with the transition from intermediate to clastic stage but appeared to decrease slowly through the clastic stage (Fig. 1). It was not possible to accurately measure cytoplasmic mHtt^ex1^-GFP at very late stages due to the emergence of dispersed inclusions and the large size of the clastic inclusion itself (Supplementary Fig. 4a).

### Structural changes decrease the solubility of inclusion contents

The hallmarks of inclusion transformation imply a loss of liquid or gel-like structure: the increasingly uneven, static distribution of inclusion contents and the irregularity of external shape. To probe the solubility of intermediate and clastic inclusions, we used auxin-mediated degradation to remove the labile cytoplasmic pool of mHtt-GFP.

We have found that ovoid IBs release mHtt-GFP back to the cytoplasm indicating that, under normal conditions, the ovoid IB grows because the rate of addition of material to the inclusion exceeds the rate of release.^10^ Concordantly, inclusions shrink when cytoplasmic mHtt-GFP is reduced below a threshold concentration using auxin-mediated degradation.^22^ Using the same strategy, we tested the ability of later stage inclusions to release material at low cytoplasmic mHtt-GFP concentrations by expressing mHtt-degron-GFP in cells co-expressing the plant E3 ligase, Tir1. In the presence of the synthetic auxin naphthalene acetic acid (NAA), cytoplasmic mHtt- degron-GFP dropped to background levels in many cells (Fig. 4).

**Figure 4.**
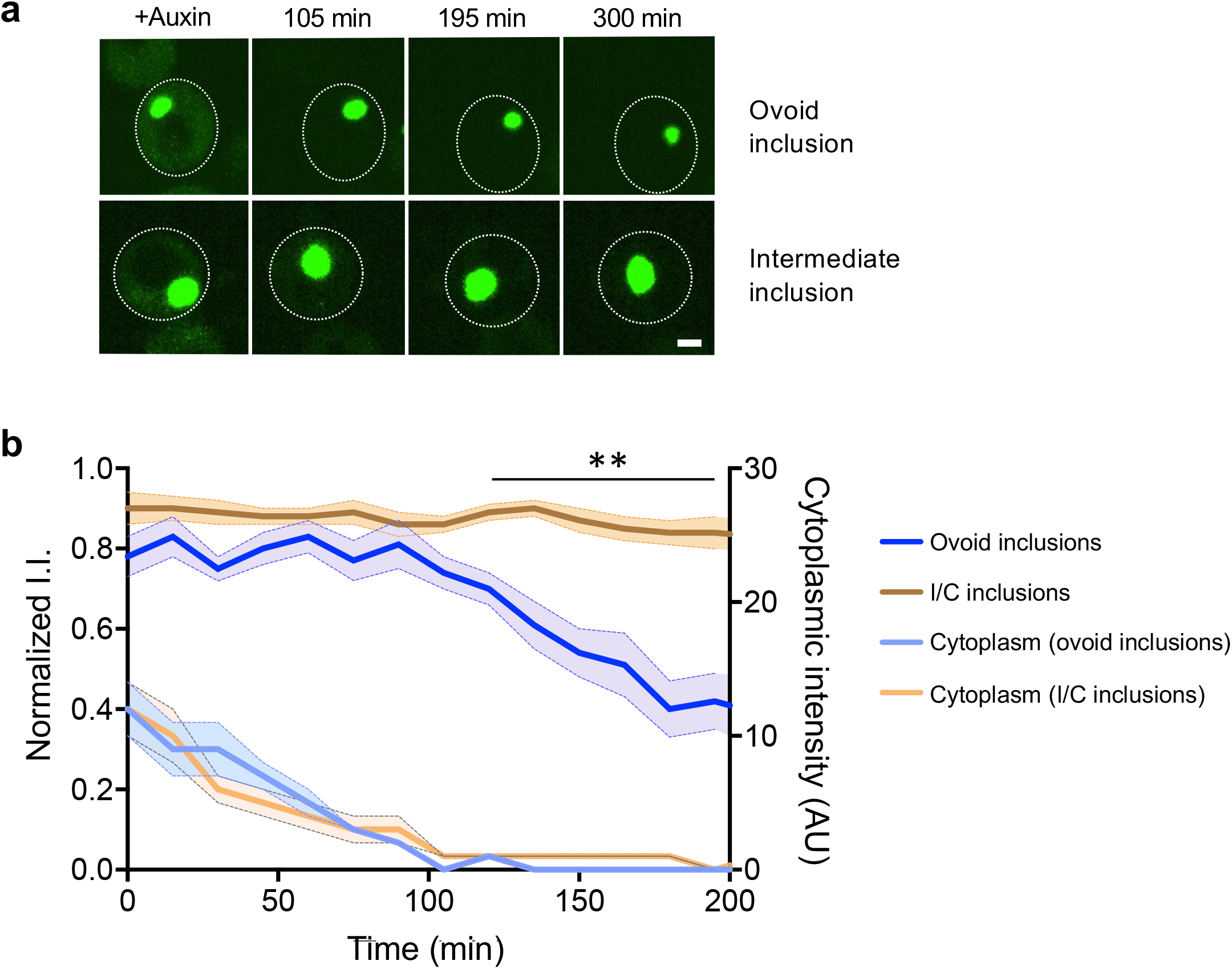
Intermediate and clastic-stage inclusions do not release material to the cytoplasm. **a**, Representative images of Htt(72Q)-degron-GFP co-expressed in cells along with E3 ligase Tir1 at the indicated times after the addition of auxin. Upper row, cell with ovoid inclusion; lower row, cell containing an intermediate inclusion. Images are identically contrast-adjusted to show the cytoplasmic intensity. Scale bar, 1 μm. **b**, Quantification of intensity of cytoplasmic Htt(72Q)- degron-GFP over time after the addition of auxin is shown, along with the integrated intensity (I.I.) of ovoid or intermediate stage and clastic-stage inclusions (I/C). The integrated intensity of individual inclusions was normalized to the highest value for that inclusion; curves were aligned at the point when the cytoplasmic intensity reached background levels (shown at 105 minutes). Lines represent the mean; shaded areas represent SEM (n=8-10 for intermediate and clastic-stage inclusions, 12-13 for ovoid inclusions). The difference between the I.I. of ovoid and I/C inclusions was significant beginning at 120 minutes (**, p≤ 0.001, multiple T-tests using the two-stage linear step-up procedure of Benjamini, Krieger and Yekutieli, with Q = 1%. Consistent SD was not assumed).

We sorted inclusions in these cells into ovoid or intermediate/clastic (late-stage) categories. The late-stage inclusions included both intermediate and early clastic inclusions for which cytoplasmic mHtt was still visible in the initial image immediately after the addition of auxin. We cannot rule out the possibility that part of the drop in cytoplasmic intensity was due to the natural disappearance of cytoplasmic mHtt during the transition from ovoid to late-stage inclusions; however, the dynamics of the decrease in cytoplasmic intensity were similar in cells containing ovoid and late-stage inclusions (Fig. 4b). Additionally, cytoplasmic intensity became undetectable in most cells with intermediate or early clastic stage inclusions, earlier than in the untreated cells we observed.

Our results show that late-stage inclusions, unlike ovoid inclusions, do not support bidirectional flux of protein between the cytoplasm and inclusion. Once the cytoplasmic protein was degraded, new protein could no longer be added to the inclusion. Ovoid inclusions shrank when cytoplasmic protein levels become very low, whereas late-stage inclusions did not shrink, demonstrating that they do not release material back to the cytoplasm at a detectable rate (Fig. 4b).

### Structural changes in inclusions lead to toxicity

Inclusion transformation led to an abundance of insoluble material inside the cell. Previous work demonstrated that expression of mHtt-GFP had little or no effect on culture growth,^23,24^ suggesting that mutant Htt-GFP is not toxic in yeast. However, insoluble inclusions are toxic in mammalian cells^15^ and bulk culture growth measurements may not be sensitive enough to detect effects, such as premature senescence, occurring in a small percentage of cells. Direct measurement of the mean generation time (MGT) of individual cells has been shown to be a powerful method of detecting entry into senescence, because senescence and death are preceded by a significant lengthening of the cell cycle.^25,26^

We used time-lapse imaging to observe divisions in cells containing a range of inclusion sizes and types (Fig. 5a). We identified 40 cells that contained clastic inclusions for the duration of the timecourse, and an additional 30 inclusion-containing cells that spent approximately half the imaging timecourse in the intermediate or clastic stage, or both. Our control cells began the timecourse with a small ovoid IB or no inclusion (67 cells). To control for differences in temperature between timecourses, the MGT of cells with late-stage inclusions was normalized to that of cells with ovoid or no inclusions for each imaging timecourse separately. The MGT of cells containing clastic inclusions was almost 3-fold higher than for cells carrying ovoid or no inclusions, and cells with intermediate inclusions had a 1.7-fold increase in MGT compared to cells with ovoid or no inclusions (Fig. 5a, b).

**Figure 5.**
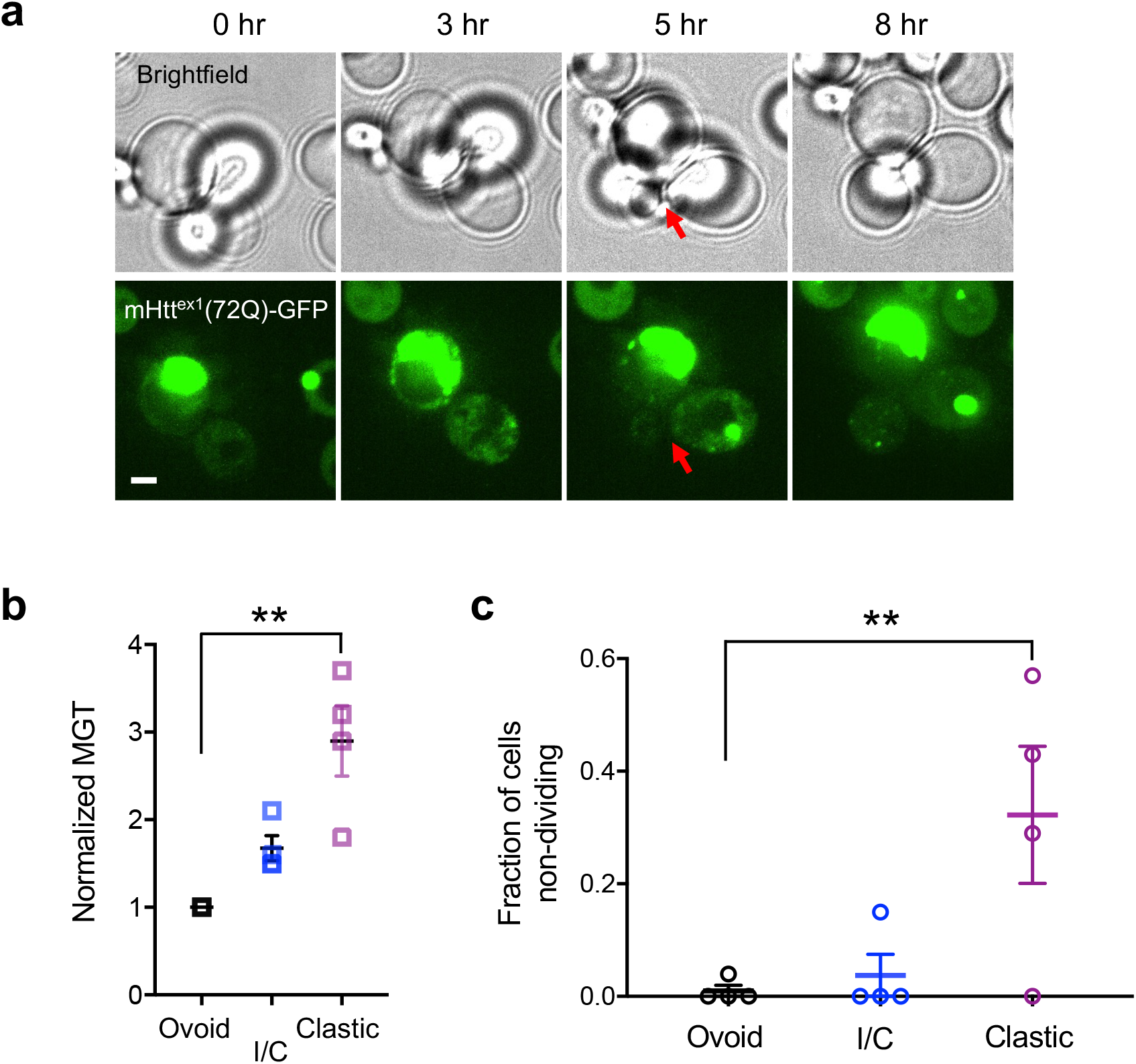
Large insoluble inclusions cause senescence and death. **a**, Representative time-lapse images in brightfield (upper row) and the GFP channel (lower row) of a mHtt(72Q)-GFP-expressing cell containing a large clastic inclusion. The cell divided once during an 8-hour timecourse (arrow indicates bud neck). Scale bar, 2μm. **b**, Mean generation time (MGT) was measured for cells containing ovoid or no inclusions; clastic inclusions; and inclusions that transitioned into intermediate or later stages during the timecourse (I/C). For each imaging timecourse, MGT was normalized to the MGT for ovoid or no inclusions. Each point represents 1 timecourse; bars represent mean and SEM of all timecourses (4 individual timecourses, two of 72Q and two of 103Q; n = 5 – 25 cells per timecourse, total n = 60, 30 and 40 cells with ovoid/no, I/C, and clastic inclusions, respectively). The difference between the MGT of cells containing ovoid and clastic inclusions was highly significant (p = 0.004, Kruskal-Wallis test with Dunn’s correction for multiple comparisons). **c**, For the data set shown in **b**, the fraction of cells is shown that did not divide at all during 8 hours of observation. The difference in the fraction of non-dividing cells between cells containing ovoid inclusions and those with clastic inclusions was highly significant (p= 0.001, one way ANOVA with Tukey’s multiple comparisons test).

Of the cells containing ovoid or no inclusions, only one of 67 cells did not divide during the timecourse; the low fraction of senescing cells is consistent with previously published rates of senescence in young BY4741 cells.^27–29^ In contrast, about 30% of cells containing clastic inclusions did not divide even once during the timecourse, a strikingly high rate of senescence (Fig. 5c).

### Late-stage inclusions are found over a broad age range, including in young cells

We considered the possibility that the underlying cause of senescence in cells containing clastic inclusions was aging rather than the accumulation of unfolded protein. Because inclusions grow over time, we expected that the transformation of the inclusion from ovoid inclusion body to clastic inclusion would also correlate with cell replicative age. To test whether increased age explains the rise in MGT and senescence in cells with clastic inclusions, we stained cells with Calcofluor White (CFW) to identify bud scars, which are a measure of a cell’s replicative age. (Supplementary Fig. 5a). First, we examined the ages of randomly selected cells without regard to their inclusions. The overall distribution of cell ages seen in randomly sampled cells expressing the 25Q, 72Q, and 103Q isoforms of mHtt were similar to each other and to the age distribution of cells which were not expressing mHtt-GFP (Supplementary Fig. 5b).^30^ Next, we examined the likelihood of cells containing inclusions at different ages. Overall inclusion frequency increased with cell age and polyQ tract length: 100% of 72Q-expressing cells with a replicative age > 6 and 103Q-expressing cells aged > 3 contained inclusions (Supplementary Fig. 5c). Finally, we ascertained inclusion stage in randomly surveyed cells. The fraction of cells containing intermediate, clastic or dispersed inclusions increased with age (Supplementary Fig. 5d), although the distribution of clastic inclusions in young cells appears bimodal; we believe that the bimodality is in part due to inclusion inheritance, and in part to differences in mHtt expression level.

To complement our randomly sampled data set, we determined the age structure of cells containing intermediate and clastic inclusions. Of cells that contain clastic inclusions, 90% of the cells were young (Fig. 6a, b). Even though middle-aged cells were increasingly likely to contain intermediate or clastic inclusions, the chance of selecting a cell that was ≥ 7 divisions old was small compared with the chance of selecting a younger cell that had a lower probability of containing a large inclusion but was more frequently represented in the overall population. We conclude that cells containing clastic inclusions were senescing at an unusually early age.

**Figure 6.**
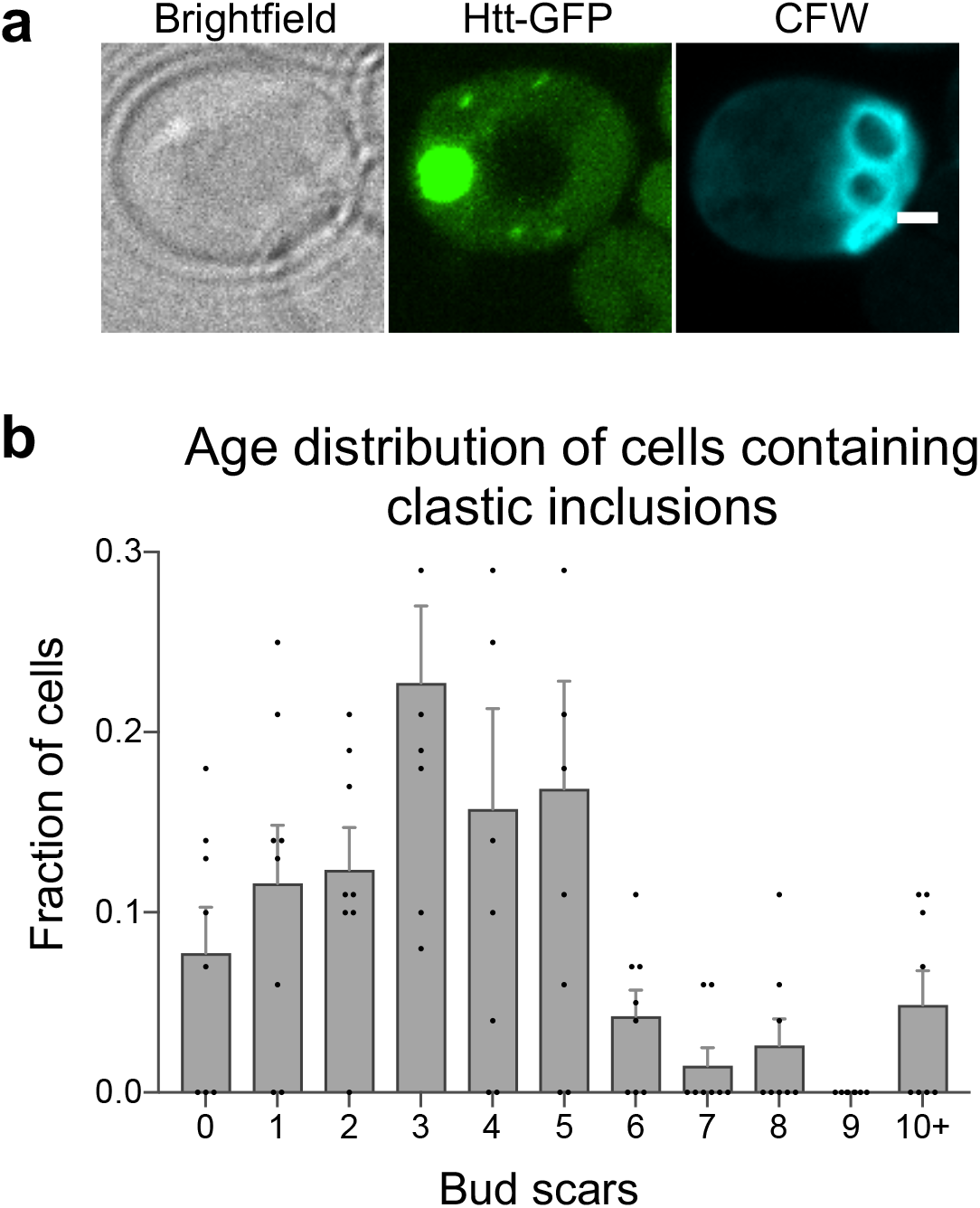
Clastic inclusions are found in young cells. **a**, Maximum projection of a cell with a clastic inclusion stained with calcofluor white (CFW). The CFW image has been contrast-adjusted to show the bud scars clearly; the bud was faintly stained by CFW but is not visible here. Scale bar, 1 μm. **b**, Age distribution of cells selected for the presence of a clastic inclusion. Four replicates each of 72Q and 103Q are shown (error bars represent SEM; n = 8 individual data sets containing 9-40 cells each, a total of 140 cells).

### Loss of mitochondrial membrane potential and cell death are correlated with late-stage inclusions

In yeast, the length of post-mitotic viability has been shown to be inversely proportional to the reproductive age at senescence: early senescence appears to lead to lengthy post-mitotic lifespans.^27^ To assess the effects of mHtt-GFP on cell viability, we stained cells with the membrane-impermeant dye propidium iodide (PI). PI enters cells with compromised membrane integrity; most cells which accumulate PI are dead or will die imminently.^31^ Strains expressing mHtt(25Q)^ex1^-GFP, 72Q and 103Q and stained with PI were imaged and scored for incorporation of PI and for mHtt-GFP inclusion stage (Fig. 7a). The 25Q-expressing strain was used as the baseline to control for cell death due to plasmid loss.

**Figure 7.**
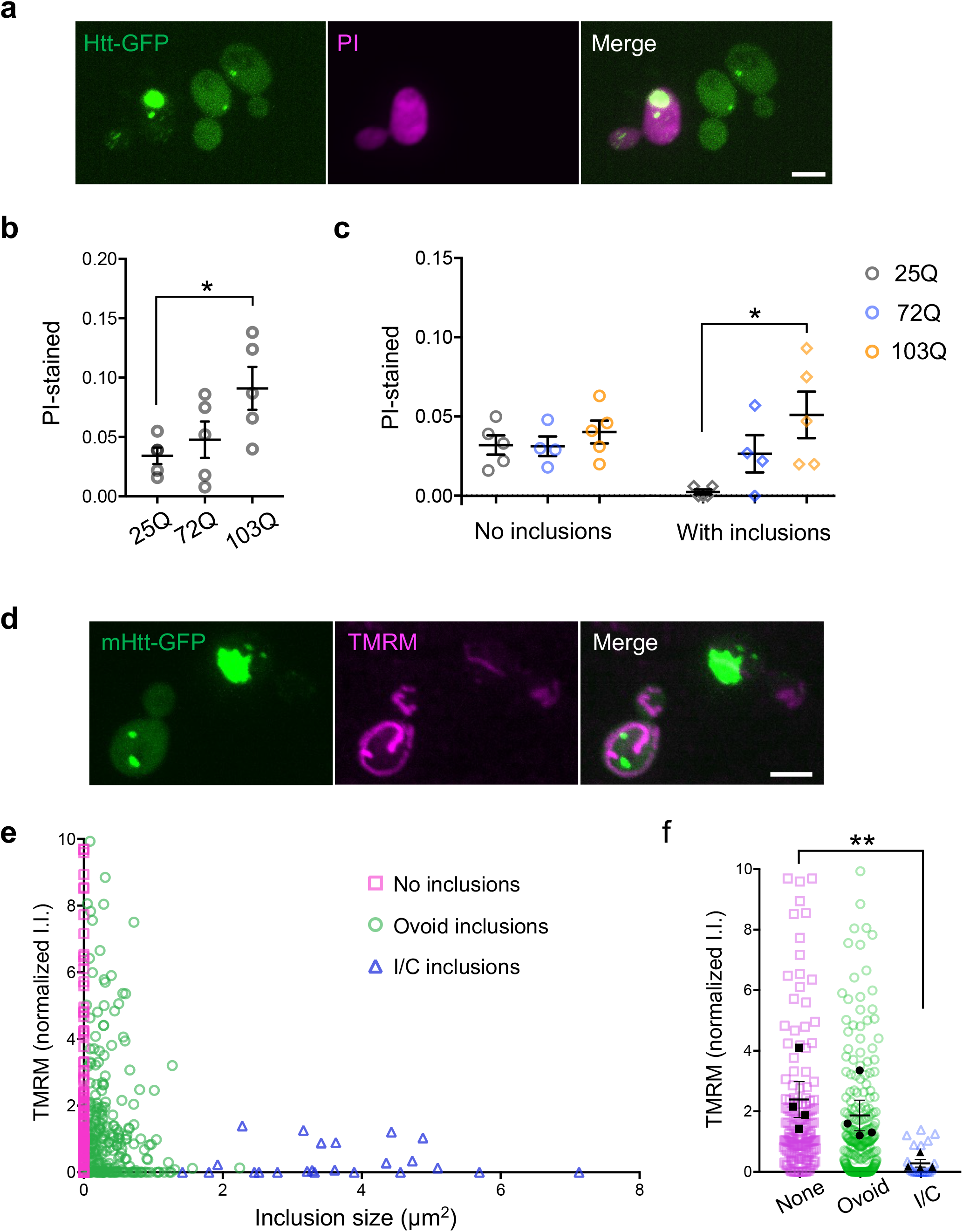
The presence of inclusions correlates with cell death and mitochondrial damage. **a**, Httex1(103Q)-GFP-expressing cells stained with propidium iodide (PI). **b**, The fraction of PI- stained cells is shown, grouped according to polyQ tract length. Each point represents the mean of a replicate (n = 72-225 cells per Htt-GFP construct per replicate, 5 replicates). There is a significant increase in the fraction of PI-stained 103Q-expressing cells compared to 25Q (p = 0.04, Kruskal-Wallis ANOVA using Dunnet’s multiple comparison’s test). **c**, Using the same data set as in **b**, the fraction of PI-stained cells with and without inclusions is shown. There is no difference in the fraction of dead cells without inclusions between Htt-GFP constructs; the increase in PI- staining 103Q cells is due to the significant increase in PI-stained cells containing inclusions compared with 25Q (p = 0.01, Kruskal-Wallis ANOVA using Dunnett’s multiple comparisons test for the comparison of PI-stained 25Q vs 103Q-expressing cells containing one or more inclusions). **d**, Httex1(103Q)-GFP-expressing cells stained with TMRM. **e**, Normalized integrated intensity (I.I.) of TMRM staining is plotted vs cross-sectional inclusion area. The y-axis is truncated at 10 in order to better show most points; the full graph is available in Supplementary Fig. 6. There is a highly significant negative correlation between λλ′_m_ and inclusion size (Spearman rank correlation = -0.29, p < 0.0001, n = 460 cells). **f**, Normalized TMRM integrated intensity (I.I.) is shown by inclusion type, showing individual points and the mean of imaging sessions (black symbols). There is a highly significant difference in λλ′_m_ between cells containing no inclusions and those containing intermediate or clastic (I/C) inclusions (p = 0.009, Friedman’s ANOVA using Dunn’s multiple comparisons test; 4 individual imaging sessions, total n = 170, 267 and 23 cells, respectively). Scale bars, 3 μm. Error bars represent the mean and SEM in **b, c** and **f**.

The frequency of PI-stained cells in the 72Q-expressing strain was similar to that seen in 25Q- expressing cells, while the fraction of PI-stained cells in 103Q-expressing cultures was elevated (Fig. 7b). The difference in PI staining frequency between 103Q and 25Q was attributable to the increased number of cells containing mHtt-GFP inclusions (Fig. 7c). Thus, the presence of inclusions is associated with loss of membrane integrity, and presumably with cell death.

Many chaperone proteins use ATP in the process of disaggregating and refolding protein, suggesting that mitochondrial function would be especially critical in cells with substantial burdens of unfolded protein. Therefore, we assessed the health of mitochondria in cells containing late-stage inclusions. Previous work by Ferhmann and colleagues^32^ demonstrated that the mitochondrial membrane potential (ΔΨ_m_) is constant over the first 12 or more divisions. Of cells containing late-stage inclusions in our studies, 98% have divided ≤ 12 times and are therefore not predicted to be undergoing age-related declines in ΔΨ_m_.

Mitochondrial membrane potential was assessed by staining with the potential-sensitive dye tetramethylrhodamine methyl ester (TMRM) (Fig. 7d, Supplementary Fig. 6a, b).^33–35^ We found that ΔΨ_m_ fell as inclusion size increased (Fig. 7e, Supplementary Fig. 6c). Mitochondrial function in cells containing late-stage inclusions was profoundly impaired in many cells: mean ΔΨ_m_ fell by 88% compared to cells with no inclusions (Fig. 7e, f). However, structural change did not require loss of ΔΨ_m_, as about a quarter of cells with late-stage inclusions had a ΔΨ_m_ close to the median value. Cellular stress and loss of ΔΨ_m_ can lead to mitochondrial fission or aggregation.^36–38^ Some cells had evidence of minor mitochondrial fragmentation, but there was no apparent correlation with inclusion state, nor evidence of mitochondrial aggregation (Supplementary Fig. 7).

### Daughter cells reverse prion-like changes, restoring soluble protein

While observing the division of cells with clastic inclusions, we noted that the loss of soluble protein was observed in the bud and was always inherited by newborn daughter cells (n = 20). In addition to the absence of diffuse mHtt-GFP, the daughter cells often contained multiple small to moderate-sized irregular inclusions (Fig. 8a, b; Supplementary Fig. 8). However, most daughter cells reversed this state, recovering soluble cytoplasmic mHtt-GFP and fusing small inclusions to form one or more ovoid inclusions. Of 18 daughter cells that we were able to observe for ³4 hours, 4 showed no recovery of either the diffuse cytoplasmic pool or an ovoid inclusion, 1 showed a partial recovery, and 13 recovered both phenotypes.

**Figure 8.**
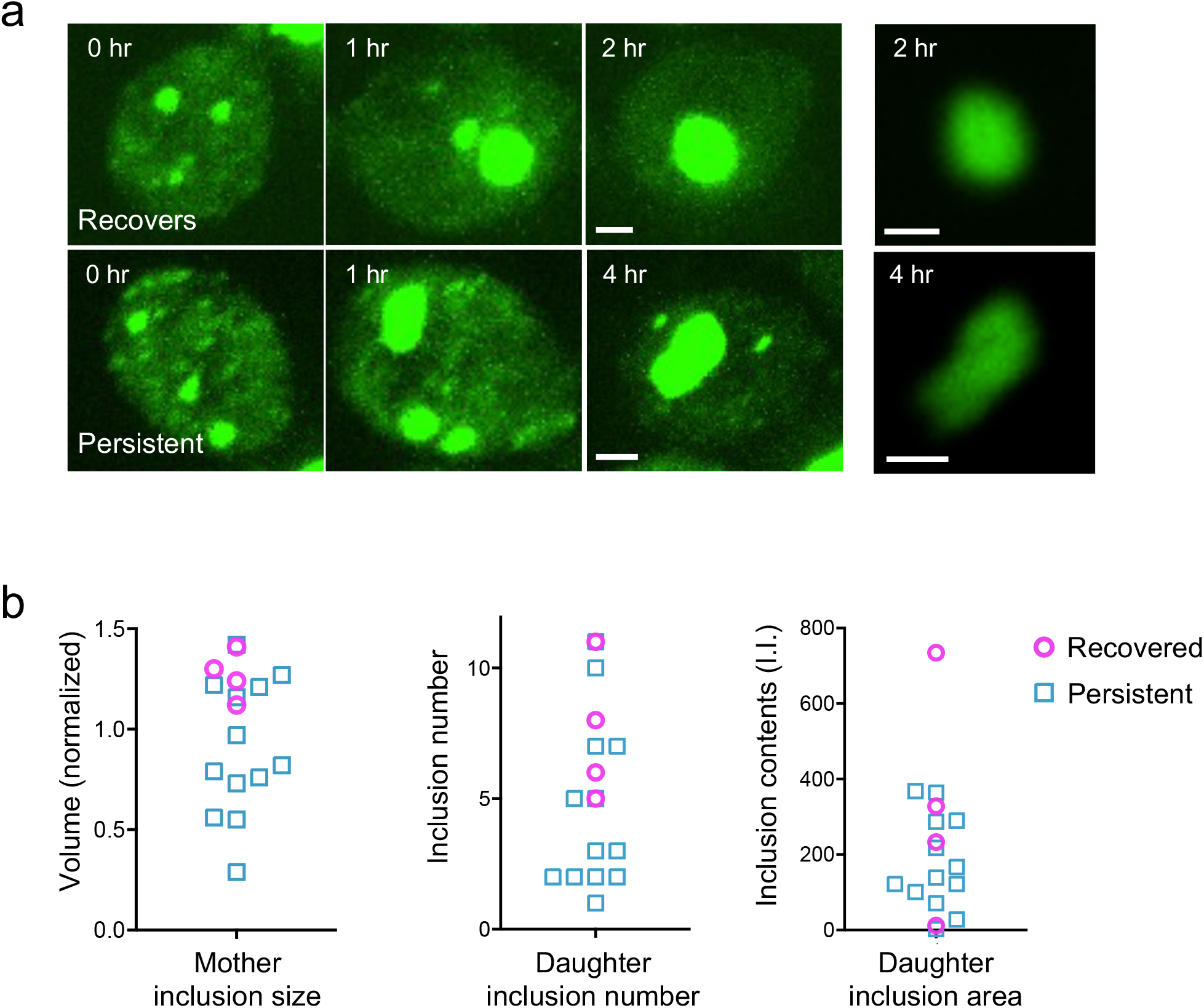
Loss of soluble mHtt protein is reversible in daughter cells. **a**, Maximum projections of two mHtt^ex1^(103Q)-GFP-expressing daughter cells at the indicated times after cell division. Upper row, a daughter cell that recovered diffuse cytoplasmic mHtt-GFP and a single ovoid inclusion; lower row, a daughter cell that did not recover. Right panels, images of the inclusion at 4 hr, contrast-adjusted to show internal distribution of mHtt-GFP. Scale bar, 1 μm. **b**, Normalized volume of the mother cell inclusion immediately before division, the number and summed integrated intensity of small inclusions in the bud immediately after division are shown, for daughter cells that recover soluble mHtt-GFP (n = 13) and daughters that do not (n = 4), in cells expressing 72Q or 103Q. There is a significant difference in mother cell inclusion volume between daughter cells that recover vs those that do not (p = 0.006, Welch’s t test). Mother cell inclusion volume was normalized within each polyQ tract length.

To understand which factors influence the reversion of inclusion stage in daughter cells, we compared mother cell inclusion size and the number and integrated intensity of daughter cell inclusions in cells that recovered, versus cells with persistent insoluble mHtt-GFP. Inclusion volume in the mother cell was the only variable that significantly correlated with failure of a daughter cell to recover (Fig. 8c). Nonetheless, mother cell inclusion size was not fully predictive, as there were many mother cells with large inclusions that gave rise to daughter cells which were able to revert to an earlier inclusion stage. Taken together, these results indicate that most daughters of cells containing clastic inclusions can reverse the prion-like state, but they are less likely to do so as the mother cell inclusion becomes larger.

## DISCUSSION

Previously, we have found that the mHtt-GFP inclusion is a condensate: it is a mobile, phase-separated cellular compartment that sequesters unfolded mHtt-GFP along with the disaggregase Hsp104 and releases mHtt-GFP back to the cytoplasm. The formation of the inclusion is promoted and regulated by the cell. Here, we show that as the inclusion expands, it undergoes a structural change concomitant with a rapid prion-like conversion of the cytoplasmic protein from a soluble to an insoluble form. This structural change is accompanied by rapid growth of the inclusion, along with appearance of smaller, dispersed inclusions, and is followed by senescence and death.

The change in inclusion structure from ovoid to clastic, like the loss of diffuse cytoplasmic protein, is consistent with a relatively rapid change from soluble to insoluble protein. In an ovoid inclusion, newly entered material diffuses throughout and the contents appear evenly distributed. In contrast, the contents of late-stage inclusions were heterogeneously distributed, with a wrinkled internal structure suggesting solidification. The distribution of another inclusion component, Hsp104- mCherry, in late-stage inclusions was identical to that of mHtt-GFP (manuscript in preparation), suggesting that this heterogeneous distribution is shared by other inclusion components and represents a global structural change. Furthermore, ovoid inclusions remained rounded as they grew, in accordance with presence of a significant surface tension, while clastic inclusions became increasingly asymmetric and convoluted, suggesting relative solidity.

In models of prion-like propagation, the seeding of cells with fibrils of an aggregative protein such as α-synuclein, SOD1, TDP-43 or Sup35 results in the loss of soluble protein and appearance of numerous inclusions.^16–18,39^ A seed, or ‘propagon’, may be transmitted to other cells, including through inheritance.^42^ In contrast, unfolded protein inclusions are normally retained in the mother during division, thus preserving the health of the daughter cell.^40,41^ An ovoid mHtt^ex1^(72Q)-GFP inclusion is retained in the mother 92% of the time (n = 85), whereas the prion-like state of mHtt-GFP was always inherited, consistent with cytoplasmic transmission of prions.

Most surprisingly, the prion-like state was reversible in young cells by a wholly physiological mechanism. Inhibition of prion oligomerization has been accomplished in vitro;^43^ enzyme inhibition, dissolution or genetic manipulations have been used to prevent prion-like transmission in yeast.^42,44–46^ Using time-lapse imaging to track inheritance of a prion-like state in individual cells, we have found that newly born cells are able to fully restore solubility both to the cytoplasmic pool and to inclusion contents. Thus, newly born cells can reverse the self-propagation of prions. To our knowledge, this is the first observation of a wholly physiological reversion of a prion-like phenotype.

What is the molecular mechanism underlying the switch to a prion-like state? Structural changes became apparent when the inclusion reached a cross-sectional area of 2 mm^2^, and this threshold size correlated with both the drop in cytoplasmic levels of mHtt-GFP (Fig. 2) and Hsp104-mCherry (manuscript in preparation) and with a profound loss of mitochondrial membrane potential (Fig. 7). These data suggest that the changes in cytoplasmic and inclusion mHtt-GFP may both be precipitated by an inclusion size threshold.

Because the first evidence we see of structural change is a ‘hollowing out’ (increase in void area), the process is reminiscent of the structural collapse of gels in vitro,^47^ which would explain the temporary plateau in inclusion growth as the shift to a denser structure temporarily offsets the continued accumulation of new material. Loss of ΔΨ_m_ was apparent around the same time as the structural transition, suggesting that a decrease in ATP, either as an enzyme substrate or as a hydrotrope,^48^ could be important in triggering the observed changes.

Contraction and hardening of a gel may be explained by a process such as syneresis, in which liquid is excluded and the gel structure becomes more compact.^49,50^ Alternatively, an increase in the concentration or volume fraction of a constituent component of a mixed colloidal-polymer gel can also precipitate a phase change to a more solid state.^51^ Ovoid inclusions are likely to contain particles containing amyloid fibers of Htt-GFP together with Hsp104.^10^ An increase in mHtt-GFP amyloid fibers relative to other inclusion components could explain the shift to prion-like propagation and inclusion solidification.

The shift in cellular mHtt-GFP solubility at a threshold inclusion size, leading to cytotoxicity, may have direct implications for therapeutic approaches, as could elucidation of the molecular mechanism underlying the physiological reversal of a prion-like state.

## MATERIALS AND METHODS

### Strains, plasmids and growth conditions

All strains used in this study were derived from BY4741 and are described in detail in Supplementary Table 1. Cultures were grown using the appropriate selective minimal media at 30°C. Transformations were performed using a standard lithium acetate protocol.^52^ PCR amplifications used a high-fidelity polymerase and amplified coding regions were sequenced before use. Plasmids used in this study are described in Supplementary Table 2.

### Confocal image acquisition

Imaging was performed using a 100x/1.45 CFI Plan Apo Lambda objective lens on a TiE2-PFS microscope (Nikon, Melville, NY) equipped with a CSU-X1 spinning-disk unit (Yokogawa Electric, Tokyo, Japan), a Zyla sCMOS camera (Andor, Belfast, Northern Ireland) and OBIS LX 488 and LS 561 lasers (Coherent Inc, Santa Clara, CA). Z-stacks were acquired with 200 nm spacing. Exposure times and laser power were kept consistent between strains, as well as between imaging sessions.

Cultures were inoculated at low density and grown overnight to mid-log phase at 30°C with shaking. For time-lapse experiments, cells were transferred to a glass-bottom dish (Mat-Tek, Ashland, MA) as previously described.^22^ Fields of cells containing one or more cells with an intermediate, clastic or large ovoid inclusion were selected and imaged once per hour for 6-10 hours. We have previously observed that the rapid turnover of mHtt-GFP prevents fully correcting mHtt-GFP images for photobleaching. Therefore, we spaced out imaging timepoints to minimize photobleaching and did not perform intensity correction.

### Auxin treatment

Auxin treatment and image analysis were performed as previously described.^22^ In brief, cells were transferred to a glass-bottom dish and 8-12 fields containing at least one cell with a late-stage inclusion were chosen for imaging. Naphthalene acetic acid (Sigma Aldrich) was added to the medium to a final concentration of 250 mM, and z-stacks were collected every 15 minutes for 4-5 hours.

### Staining

#### Bud scar staining

Bud scars were stained using calcofluor white (CFW, Millipore Sigma). Mid-log phase cells were resuspended in 0.25x PBS containing 5% glucose and CFW was added to 2 mg/mL. The cells were incubated for 1 minute at room temperature in the dark with shaking. Cells were pelleted by centrifugation at 1800xg for 3 minutes and resuspended in 0.25x PBS containing 5% glucose and transferred to a slide for live imaging. To determine bud scar distribution, fields of cells were chosen in bright-field to ensure random selection. For selective analysis of cells containing transitional and late-stage inclusions, cells were scanned in the green channel and fields containing cells with late-stage inclusions were imaged. Bud scars were counted manually by visualization with the ImageJ 3D Viewer plugin.^53^

#### Determination of cell death

Mid-log phase cells were resuspended in 0.25x PBS containing 5% glucose and propidium iodide (PI) was added to 5 mg/mL. Cells were incubated with rocking for 10 minutes, pelleted, resuspended in 0.25x PBS containing 5% glucose as above, and transferred to slides for imaging. Cells were imaged for no more than 10 minutes. Fields of cells were chosen in bright-field to ensure random selection.

#### Determination of mitochondrial membrane potential

Tetramethylrhodamine methyl ester (TMRM) staining was performed identically to PI staining, using a final concentration of 50 nM TMRM. To confirm that staining was dependent on ′Ι′_m_, cells were pretreated with carbonyl cyanide m-chlorophenyl hydrazone (CCCP) in buffer at a final concentration of 25 μM for 10 minutes prior to the addition of TMRM. Cells were imaged for no more than 10 minutes. Images were analyzed as described below.

#### Image processing and analysis

Image analysis was performed on unprocessed images using the Fiji distribution of ImageJ.^19^ Statistical analyses were performed in Microsoft Excel or GraphPad Prism 7. Scripts are available at https://github.com/theresaswayne/Htt_Inclusions.

#### Determination of inclusion stage, area and cytoplasmic intensity

Cytosolic intensity was measured in a representative region of the cell, excluding the vacuole and distant from the inclusion. A threshold of 1.5x the mean cytosolic intensity was used to identify the inclusion/s; if dispersed inclusions were present, the large inclusion was manually selected for analysis. The optical section of highest intensity was assumed to approximate the central plane and was used to measure cross-sectional area.

Using the plane of highest intensity, the inclusion image was segmented using Niblack’s auto local thresholding algorithm.^21^ The binary processed image was used to measure the number of independent objects and their areas. The ImageJ ‘Fill Holes’ function was applied, and the number of independent objects and their areas were measured again. If multiple areas were reported after local thresholding, the inclusion was categorized as clastic. If a single area was reported, the void area was calculated as 1 – (area before filling / area after filling).

#### Quantitation of inclusion volume and surface area

The cytoplasmic intensity outside the inclusion was measured and a threshold equal to 2x the cytoplasmic intensity was set to reduce artifacts generated by blur at the top and bottom of the inclusion. The Fiji 3D Objects Counter was used to measure the volume and surface area of the inclusion.^54^ The absolute value of inclusion volumes was inflated due to spherical aberration; however, the axial distortion did not appear to be significantly affected by inclusion size and therefore did not affect the relative volumes shown in Fig. 3 and Supplementary Fig. 3 and 4.

#### Measurement of mitochondrial membrane potential

TMRM was quantified using maximum projected Z stacks. A threshold of 1.2x mean cytosolic intensity was used to identify mitochondria. Area and mean intensity of pixels above threshold were used to calculate integrated intensity after correction for image background. Consistent with previous published work, TMRM mean intensity decreased during a 10-minute imaging session (Klier et al, 2021). For this reason, each image was treated as an independent data set and TMRM integrated intensity was normalized to the median for each image. Cells containing late-stage inclusions were excluded when calculating the median TMRM integrated intensity for each image.

#### Assessing mitochondrial morphology

Cells expressing mitochondrially targeted TagBFP and co-expressing either mHtt^ex1^(72Q)-GFP or mHtt^ex1^(103Q)-GFP were imaged; an experienced mitochondrial cell biologist (TS) visually assessed the blue channel for mitochondrial fragmentation or aggregation, blinded with respect to Htt-GFP (n = 104 and 131 cells respectively, including 15 cells containing late-stage inclusions).

#### Q*uantitation of Htt-GFP and Hsp104mCherry*

Htt-GFP and Hsp104-mCherry cytoplasmic intensity and integrated intensity inside the inclusion were measured using a script, previously described.^22^ In brief, mean cytosolic intensity was measured and the inclusion was segmented using a threshold of 1.5x the mean cytoplasmic intensity in the optical section with the highest maximum intensity. The Analyze Particles function was used to measure the area and mean intensity of the inclusion. Mean cytoplasmic intensity was also reported. The integrated intensity was calculated from the inclusion area and mean intensity after correction for the image background.

## ACKNOWLEDGEMENTS

The authors are grateful to Istvan Boldogh for advice and assistance with strain construction, Liza Pon and Alexander Birk for reagents, and Jeffrey Morris for valuable discussions. We would also like to thank Andrew Gunner for his contribution to preliminary data acquisition. This work was supported by grants from the National Institutes of Health to LE (SC3GM136517). The spinning-disk confocal microscope was acquired through Department of Defense Equipment Award W911NF-17-1-0516.

