## Supplemental Materials for "Threshold inclusion size triggers conversion of huntingtin to prion-like state that is reversible in newly born cells"

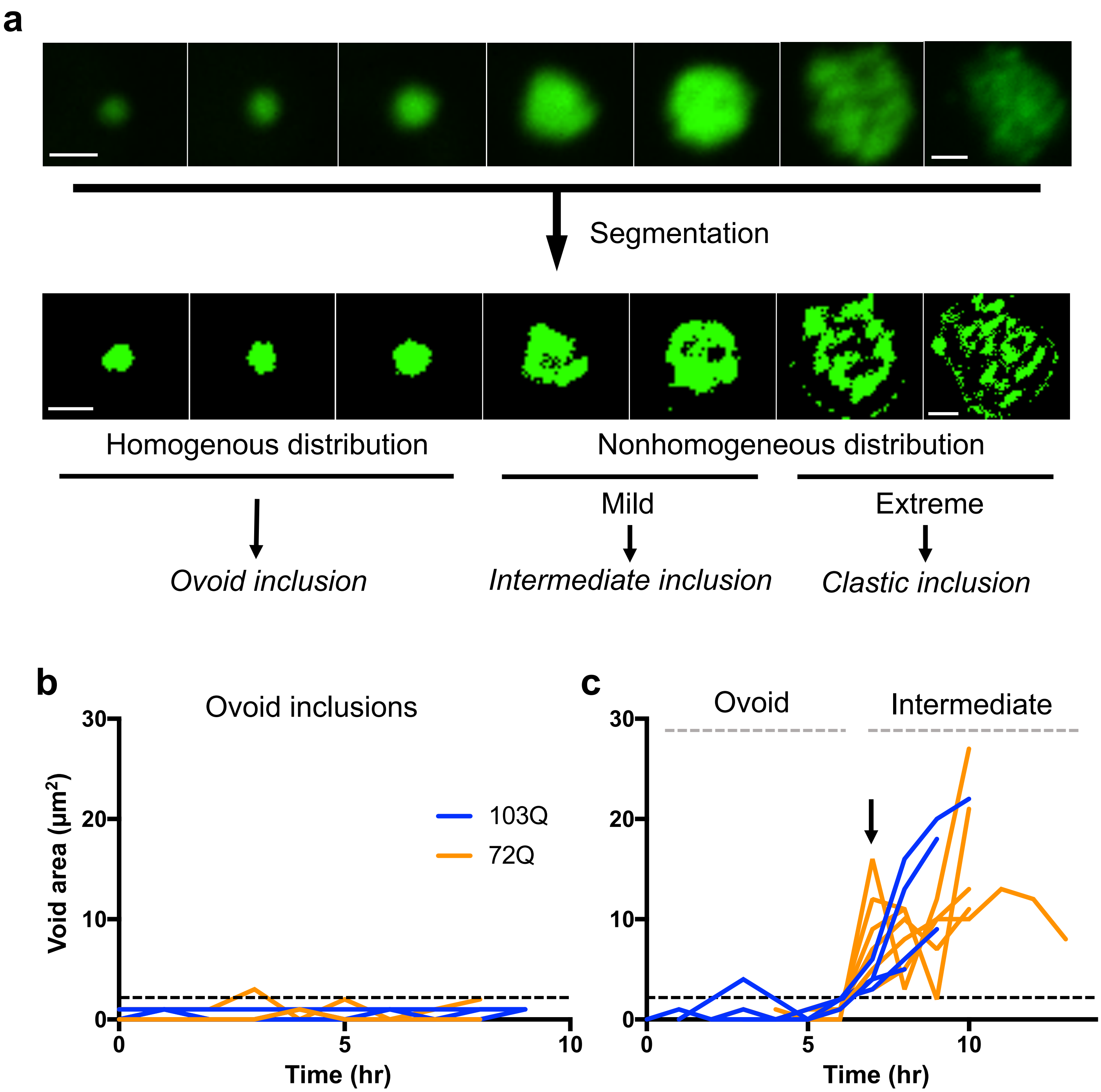


**Supplementary Figure 1. Parameters used to categorize inclusion stage. a**, Upper row: A magnified image of the inclusions shown in Figure 1. Lower row: Heterogeneities in the inclusion were identified by segmentation using the Niblack local thresholding algorithm. **b,** Void area over time for 6 ovoid inclusions, each imaged for 6-8 hours. Dashed horizontal line indicates 2%. **c,** Void areas over time for 10 individual inclusions were aligned at the point of transition (timepoint 6, indicated with an arrow). Scale bar, 1μm.

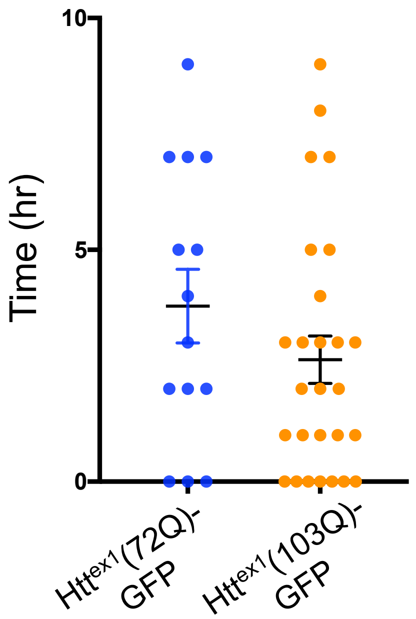


**Supplementary Figure 2. Duration of intermediate stage.** The average time that a cell contained an intermediate stage inclusion is shown separately for 72Q and 103Q-expressing cells. The difference in mean is not significant ($p = 0.23$, Welch's t test; $n = 53$, 71 cells respectively).

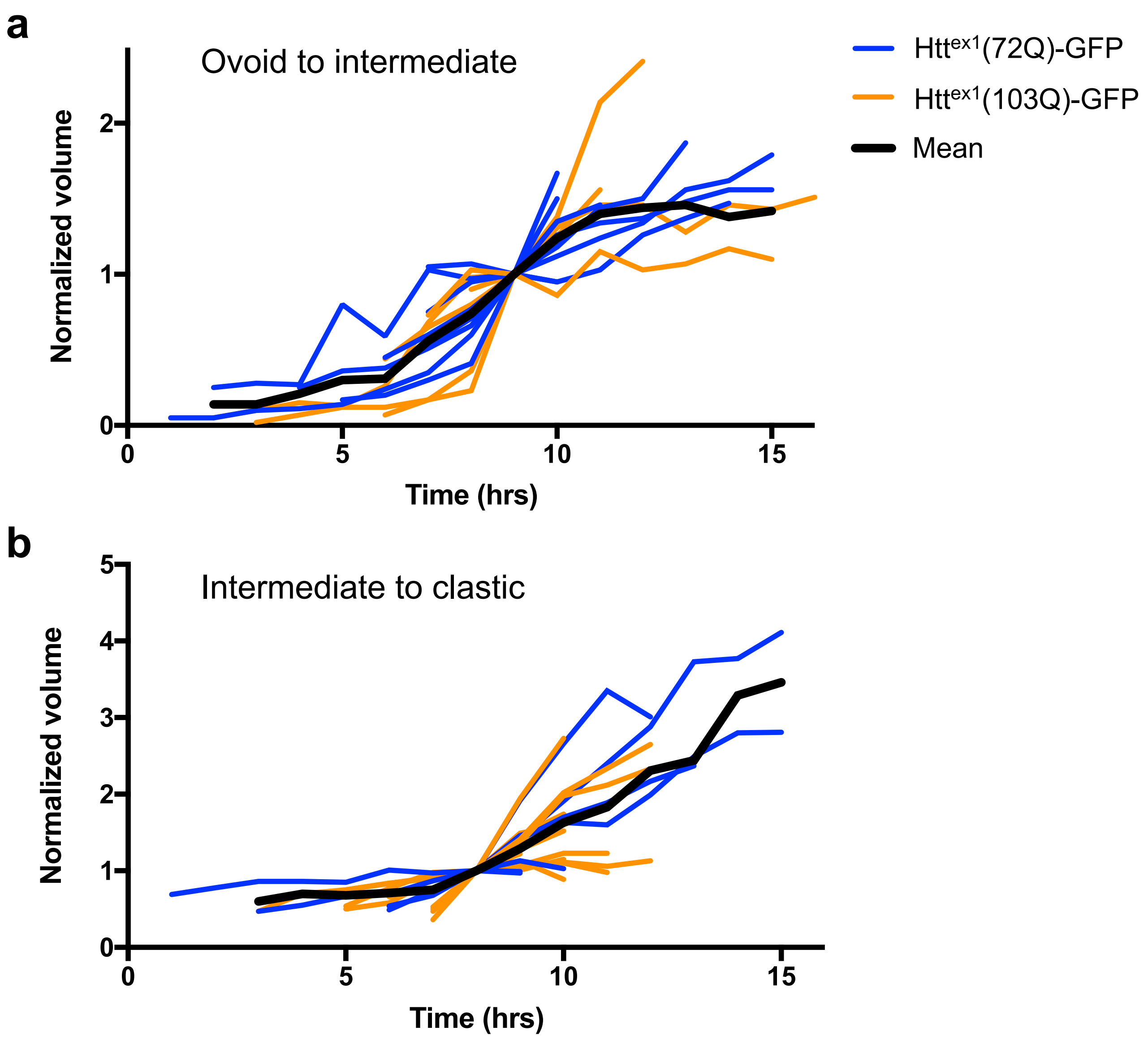


**Supplementary Figure 3. Volume of individual inclusions at each structural transition.** Inclusion volume for the transition from ovoid to intermediate, **a,** and intermediate to clastic, **b**. Blue lines are individual mHtt$^{ex1}$(72Q)-GFP inclusions, orange are mHtt$^{ex1}$(103Q)-GFP, black lines are the mean (n = 19, 18 inclusions in **a** and **b**, respectively). Volumes for each inclusion were normalized to the volume at the indicated transition, and timecourses were aligned at the point of transition.

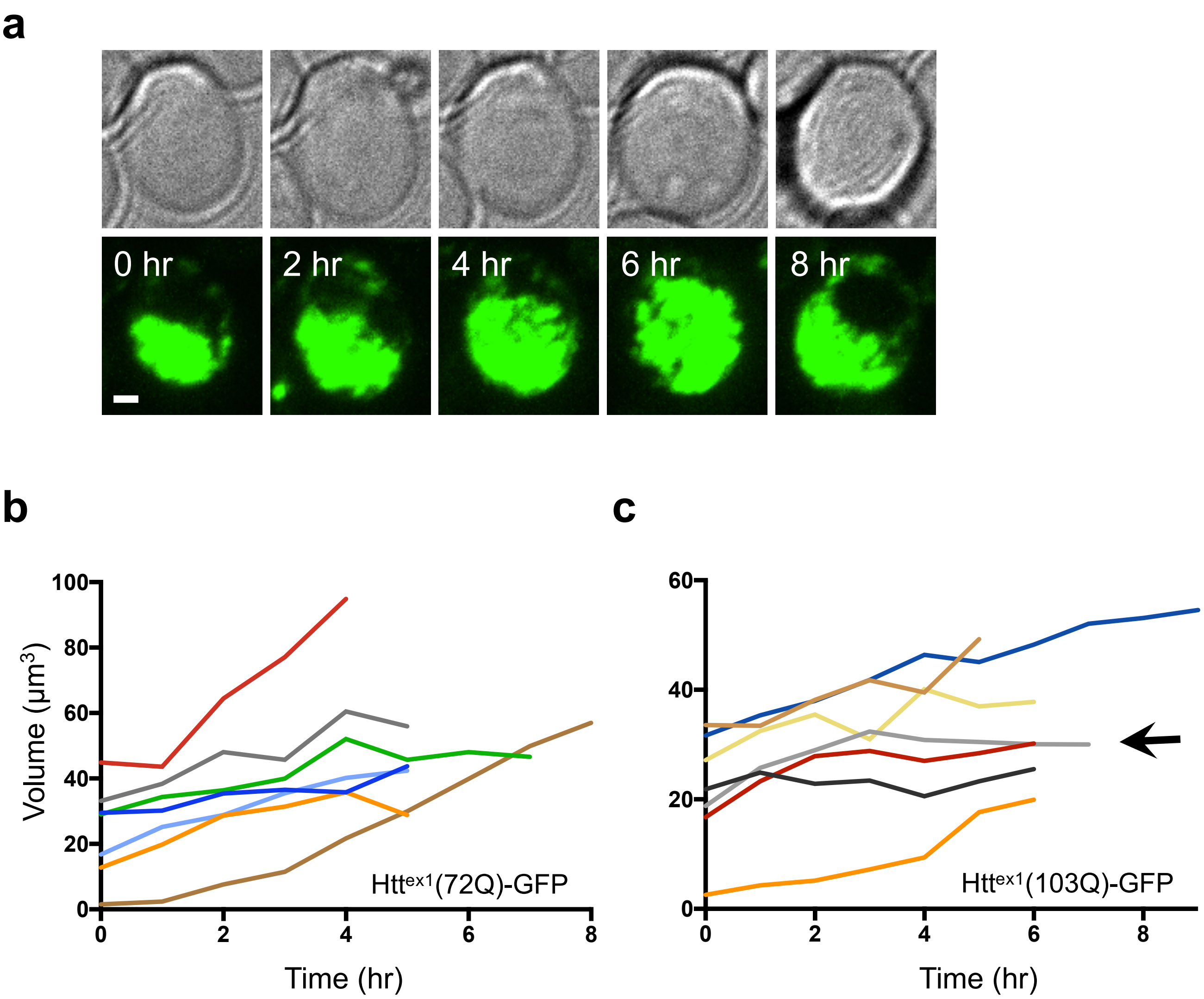


**Supplementary Figure 4. Growth of clastic inclusions. a,** Maximum projection of a representative clastic inclusion, whose growth plateaued after several hours. **b,** Measured inclusion volume for individual clastic Htt$^{ex1}$(72Q)-GFP inclusions and **c**, Htt$^{ex1}$(103Q)-GFP inclusions. Due to axial distortions, the measured volume is higher than the actual volume. The arrow in **c** indicates the inclusion seen in **a** (grey line). Inclusion volume has plateaued for some inclusions in this data set, but not all. Scale bar, 1 μm.

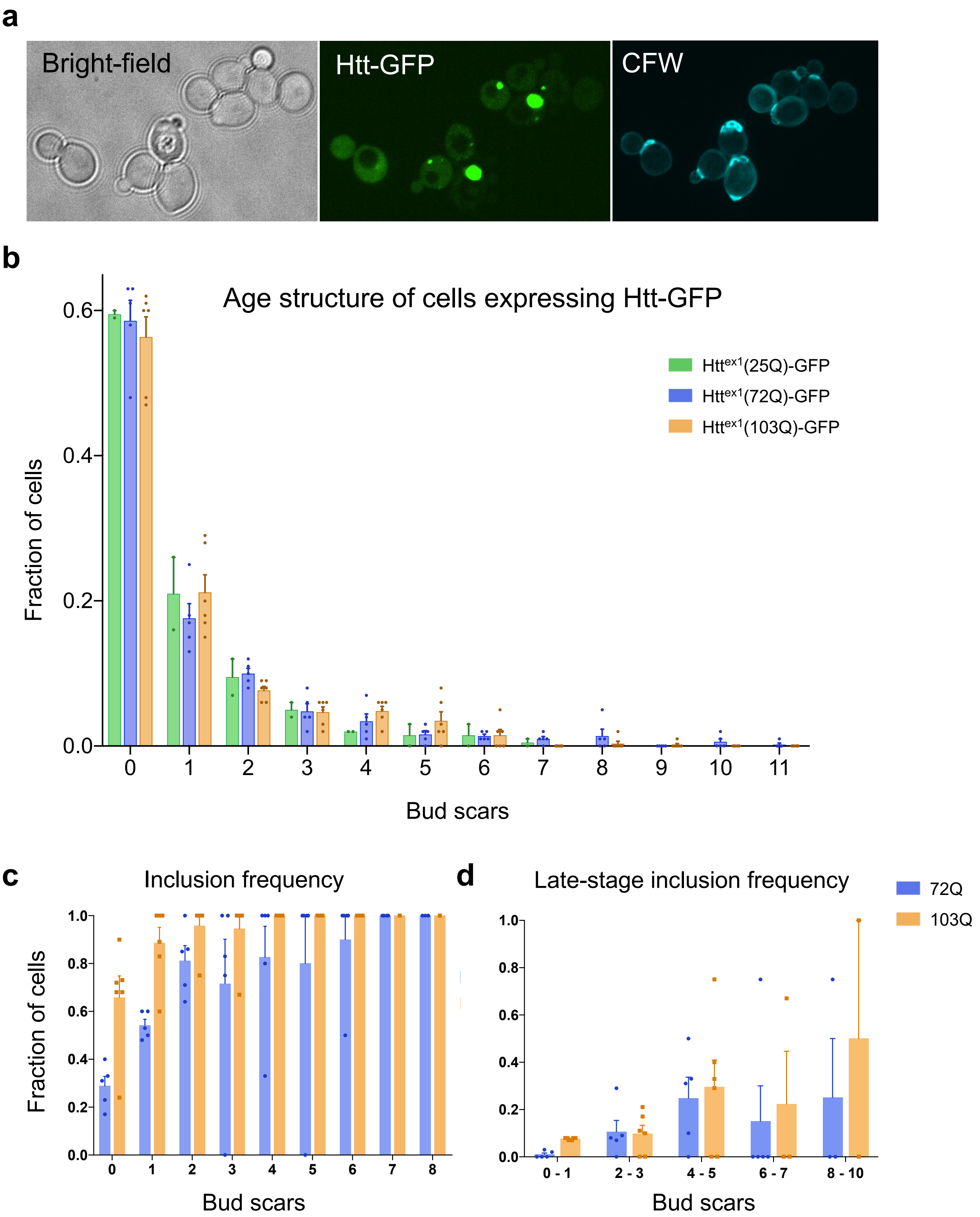

a
Bright-field
Htt-GFP
CFW
b
Age structure of cells expressing Htt-GFP
Htt^ex1(25Q)-GFP
Htt^ex1(72Q)-GFP
Htt^ex1(103Q)-GFP
Fraction of cells
0.6
0.4
0.2
0.0
0 1 2 3 4 5 6 7 8 9 10 11
Bud scars
c
Inclusion frequency
Fraction of cells
1.0
0.8
0.6
0.4
0.2
0.0
0 1 2 3 4 5 6 7 8
Bud scars
d
Late-stage inclusion frequency
72Q
103Q
1.0
0.8
0.6
0.4
0.2
0.0
0 - 1
2 - 3
4 - 5
6 - 7
8 - 10
Bud scars

**Supplementary Figure 5. Frequency of inclusions increases with age, proportional to polyQ length.** a, Maximum projection of a field of cells selected randomly in bright-field, showing mHtt-GFP (green) and Calcofluor White (CFW) staining for bud scars (cyan). **b,** The age distribution of randomly selected cells expressing 25Q, 72Q and 103Q. **c,** The fraction of randomly selected cells that contain an inclusion of any type. In **b** and **c**, bars represent the mean of 2, 5 and 6 replicates, respectively; dots represent the value of each replicate; error bars represent SEM (n = 44-125 cells per replicate, total n = 200, 465 and 465 cells, respectively). **d,** Using the same data set, the fraction of 72Q and 103Q cells containing late-stage inclusions (intermediate and clastic) at each age is shown (error bars represent SEM, ages binned to increase sample size per bin).

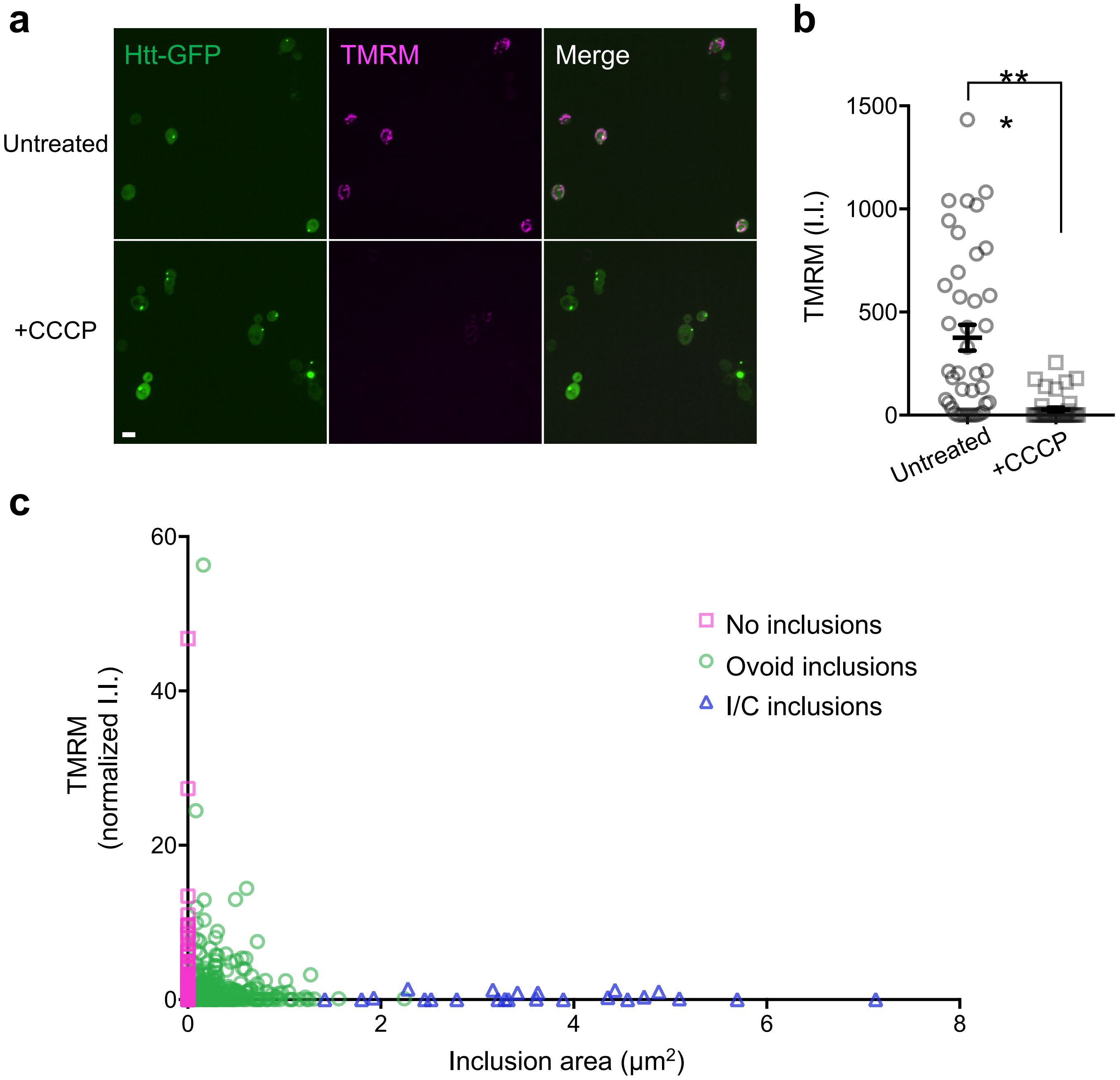


**Supplementary Figure 6. TMRM staining is responsive to mitochondrial membrane potential and decreases with inclusion size. a,** TMRM staining is lost when electron transport is blocked using carbonyl cyanide m-chlorophenyl hydrazone (CCCP). Images shown were the first obtained from fresh slides; Htt(103Q)$^{ex1}$-GFP (green) and TMRM (magenta) were contrast-adjusted using identical parameters. Scale bars, 3 μm. **b,** TMRM integrated intensity (I.I.) was compared in untreated and CCCP-treated cells to assess mitochondrial membrane potential. The first 6 images from fresh slides were quantified. The difference in integrated intensity was highly significant ($p < 0.0001$, unpaired two-tailed t test, n = 41-44 cells, respectively). **c**, Normalized integrated intensity (I.I.) of TMRM staining is plotted vs cross-sectional inclusion area showing all data points. A truncated version of this graph is shown in Figure 7.

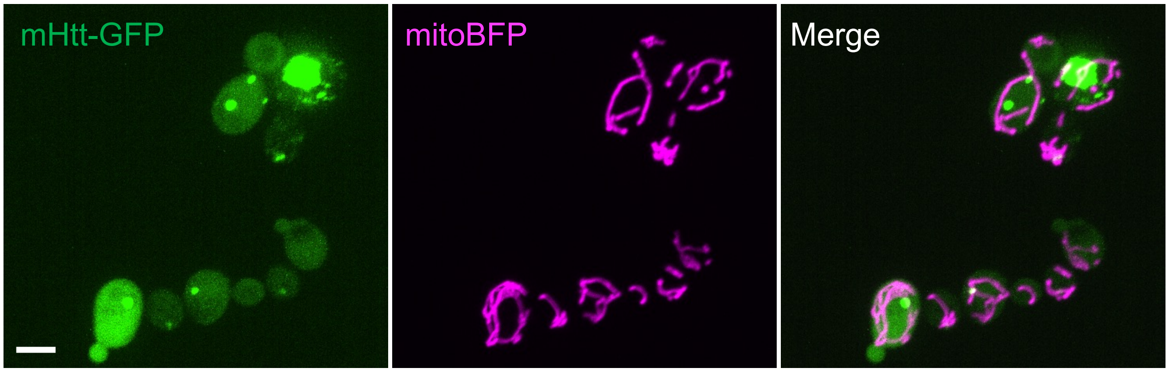


**Supplementary Figure 7. Mitochondrial morphology is not affected by inclusion state.** Representative field of cells expressing mHtt$^{ex1}$(103Q)-GFP (green) and mitochondrially targeted TagBFP (magenta). No evidence of significant mitochondrial aggregation or fragmentation was associated with late-stage inclusions (n = 235 cells, 15 containing late-stage inclusions). Scale bar, 3 μm.

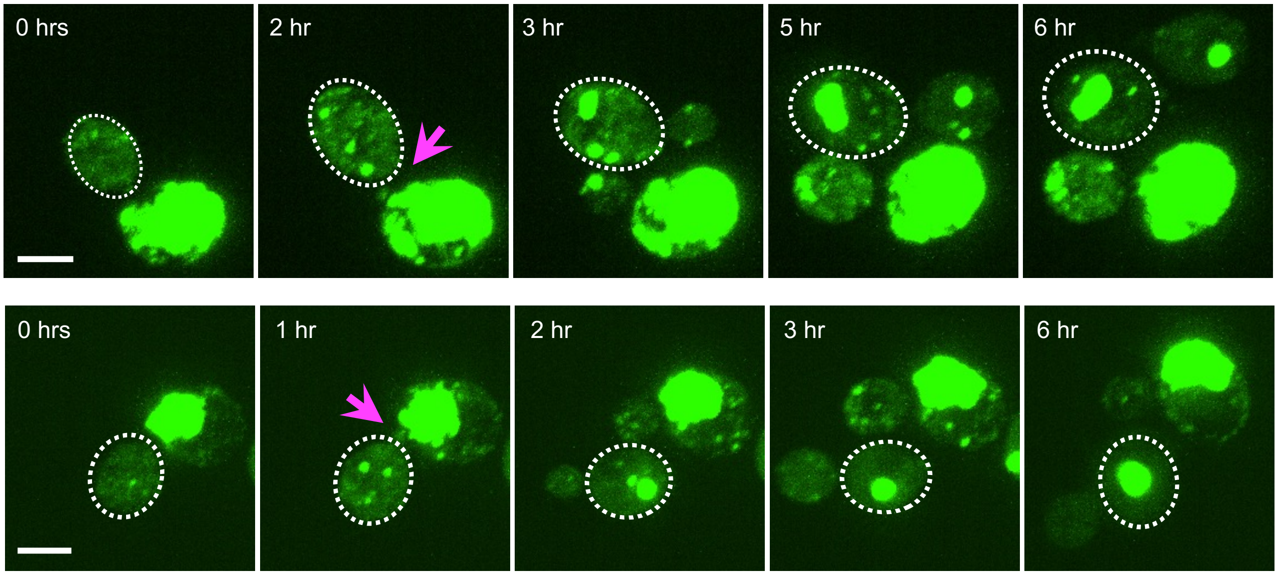


**Supplementary Figure 8. Cells with large Htt-GFP inclusions undergo prion-like loss of soluble protein that can be reversed in daughter cells.** Maximum projection of mother and daughter cells shown in Fig. 8; individual image times are indicated. Daughters are outlined; the timepoint immediately after division of the daughter cell is indicated (arrow). Scale bar, 3μm.

**Supplementary Table 1. A list of strains used in this study.**

| **Strain** | ***Genotype*** | ***Source*** |
|---|---|---|
| mHtt(72Q)-GFP | *MATa his3Δ1 leu2Δ0 met15Δ0 ura3Δ0 pEB4 [pGDP-mHtt(72Q)-GFP::LEU2]* | Aktar et al., 2019 |
| mHtt(103Q)-GFP | *MATa his3Δ1 leu2Δ0 met15Δ0 ura3Δ0 pEB11 [pGDP-mHtt(103Q)-GFP::LEU2]* | Pei et al., 2021 |
| Htt(25Q)-GFP | *MATa his3Δ1 leu2Δ0 met15Δ0 ura3Δ0 pEB18 [pGDP-mHtt(25Q)- GFP::LEU2]* | Aktar et al., 2019 |
| mHtt(72Q)-degron-GFP | *MATa his3Δ1 leu2Δ0 met15Δ0 ura3Δ0 pEB18 [pGDP-mHtt(72Q)-degron-GFP::LEU2]* | Pei et al., 2021 |
| HO-TIR1 | *MATa his3Δ1 leu2Δ0 met15Δ0 ura3Δ00 HO-TIR::kanMx* | Pei et al., 2021 |
| HO-TIR1; mHtt(72Q)-degron-GFP | *MATa his3Δ1 leu2Δ0 met15Δ0 ura3Δ00 HO-TIR1::KanMx pEB18 [pGDP-mHtt(72Q)-degron-GFP::LEU2]* | Pei et al., 2021 |
| Hsp104-mCherry; mHtt(72Q)-GFP | *MATa his3Δ1 leu2Δ0 met15Δ0 ura3Δ0 Hsp104-yEpolylinker-mCherry-hphMX4 pEB4 [pGDP-mHtt(72Q)-GFP::LEU2]* | Aktar et al., 2019 |
| Hsp104-mCherry; mHtt(103Q)-GFP | *MATa his3Δ1 leu2Δ0 met15Δ0 ura3Δ0 Hsp104-yEpolylinker-mCherry-hphMX4 pEB4 [pGDP-mHtt(103Q)-GFP::LEU2]* | This study |
| mHtt(72Q)-GFP; mitoTagBFP | *MATa his3Δ1 leu2Δ0 met15Δ0 ura3Δ0 pEB4 [pGDP-mHtt(72Q)-GFP::LEU2, pADH1-mito-TagBFP::URA3]* | This study |
| mHtt(103Q)-GFP; mitoTagBFP | *MATa his3Δ1 leu2Δ0 met15Δ0 ura3Δ0 pEB4 [pGDP-mHtt(72Q)-GFP::LEU2, pADH1-mito-TagBFP::URA3]* | This study |

**Supplementary Table 2. A list of plasmids used in this study**

| Plasmid | Description | Source |
|---|---|---|
| pEB4 | Htt(72Q)-GFP/CEN/LEU2, derived from p415GPD | Aktar et al., 2019 |
| pEB11 | Htt(103Q)-GFP/CEN/LEU2, derived from p415GPD | Aktar et al., 2019 |
| pEB18 | Htt(72Q)-IAA$^{71\text{-}114}$-GFP/CEN/LEU, derived from p415GPD | Pei et al., 2021 |
| pVT100U-mtTagBFP | mtTagBFP/2μ/URA, derived from pVT100U | Liza Pon |
